# Understanding Substrate Transport and Metabolism in *Clostridium thermocellum* to Improve Biofuel Production

**DOI:** 10.64898/2026.10.15.759513

**Authors:** Isaiah D. Richardson, Bishal Dev Sharma, Elizabeth Y. Ma, Richard S. Pilny, Daniel G. Olson

**Affiliations:** Thayer School of Engineering at Dartmouth College, Hanover, NH 03755 USA; Center for Bioenergy Innovation, Oak Ridge National Laboratory, Oak Ridge, TN 37830 USA

**Keywords:** *Clostridium thermocellum*, *Hungateiclostridium thermocellum*, *Ruminiclostridium thermocellum*, *Acetivibrio thermocellus*, alternative substrates, transport, tetrazolium dye assays, batch fermentations, degree of reduction, redox balance

## Abstract

As a cellulolytic specialist microbe, *Clostridium thermocellum* primarily ferments cellulose and its breakdown products. However, the full breadth of its substrate range remains poorly understood. In this work, we investigated the ability of *C. thermocellum* to transport and metabolize a diverse range of alternative carbon substrates. We performed tetrazolium dye assays to assess the ability of wild-type (WT) *C. thermocellum* (strain LL1004) to transport and metabolize 190 different substrates and identified 18 positive hits and 27 false positives. These findings guided the selection of 22 substrates for batch fermentations of WT on 10 g/L alternative substrate plus 1 g/L cellobiose. We identified eight substrates that were either fully or partially consumed (cellobiose, glucose, fructose, sorbitol, arbutin, turanose, pyruvate, glucose-6-phosphate), two that were interconverted (malate, fumarate), seven that may undergo conversion to unidentified fermentation products (xylose, sorbose, tagatose, galactose, mannose, rhamnose, fucose), and five that were not consumed (glycerol, acetate, lactate, propionate, xylitol). Sorbitol and pyruvate, which are more and less reduced than cellobiose, respectively, were selected for further analysis of their effects on redox balance and product formation. Batch fermentations on sorbitol and pyruvate demonstrated that ethanol production in *C. thermocellum* may be limited in part by the availability of reducing equivalents. Collectively, this work demonstrates the utility of alternative carbon substrates as a tool for perturbing metabolism, redox balance, and fermentation product distributions to further our understanding of the metabolism of anaerobic, thermophilic bacteria such as *C. thermocellum* for the production of biofuels and biochemicals of interest.

**Importance:** This work has advanced our understanding of the substrate range of *C. thermocellum* beyond its known substrates and identified several substrates including arbutin, turanose, pyruvate, and glucose-6-phosphate that *C. thermocellum* can metabolize and convert to fermentation products. This work has also demonstrated the use of alternative carbon substrates as a tool for perturbing metabolism, altering redox balance, and changing fermentation product profiles to advance our understanding of a metabolic pathway of interest, such as the cellobiose to ethanol production pathway in *C. thermocellum*. These insights can be used to inform future research aimed at engineering a strain of *C. thermocellum* for the production of cellulosic ethanol at high yield, titer, and rate. This experimental approach can be broadly applied to advance our understanding of the physiology and metabolism of other microorganisms and to engineer promising microbial candidates for the biomanufacturing of biofuels, biochemicals, or pharmaceuticals of interest.

## Introduction

Cellulosic biomass is one of the largest sources of raw material in the biosphere (Himmel et al. 2007), and on an energy basis ($/gigajoule) is a cheaper feedstock than petroleum (Lynd et al. 2017). Converting this raw material to fuels and chemicals offers substantial potential benefits including energy security, reduced greenhouse gas emissions (Field et al. 2020), and increased rural employment (Dale et al. 2014; Jordan et al. 2007). To achieve these benefits, cellulosic fuels and chemicals need to be produced at large scale, with high yield, titer and rate. However, cellulosic biomass is a complex material, and its diverse array of chemical bonds and intricate macromolecular structure make it recalcitrant to deconstruction (Lynd and Weimer 2002).

The recalcitrance of this feedstock has spurred the evolution of microorganisms specializing in cellulose deconstruction—cellulose specialists—which include anaerobic bacteria in the genera *Fibrobacter*, *Ruminococcus*, and *Clostridium* (Artzi et al. 2017) and recently reclassified *Ruminiclostridium*, *Hungateiclostridium*, and *Acetivibrio* (Yutin and Galperin 2013; Ortiz De Ora et al. 2018; Lin et al. 1994; Tindall 2019). (Note: *Clostridium thermocellum; Hungateiclostridium thermocellum; Ruminiclostridium thermocellum; Acetivibrio thermocellus* are all synonyms for the same organism. We include all of them for the sake of continuity in the scientific literature. We continue to use the original name–*C. thermocellum*–until the taxonomical changes have settled down and a new organism name becomes widely accepted by the research community). The strong native ability of these organisms to ferment cellulose makes them promising candidates for the production of fuels and chemicals (Lynd and Weimer 2002). These organisms typically have a limited range of substrates they can use for growth or fermentation, and prefer to grow on crystalline cellulose and its hydrolysis products (and only reluctantly on glucose) (Lynd and Weimer 2002). Among the anaerobic bacteria capable of solubilizing cellulose, *Clostridium thermocellum* has attracted the most attention and has been subjected to the largest number of studies. One of our long-term goals is to engineer *C. thermocellum* for the biomanufacturing of biofuels and biochemicals from cellulosic biomass at a commercial scale.

Despite potential benefits, *C. thermocellum* currently exhibits product titers that are too low for commercial applications (Dien et al. 2003). Elucidating and addressing the factors that limit high product titer in *C. thermocellum* (i.e., product tolerance (Olson et al. 2023) or redox balance (Thompson et al. 2015)) is an important step in engineering this microbe for commercial use. One approach for deciphering the factors that limit high product titer in *C. thermocellum* is to advance our understanding of its non-canonical metabolism. This can be experimentally examined using alternative carbon substrates that enter central metabolism at different locations as a tool to perturb metabolism, redox balance, and fermentation product profiles. This approach may provide key insights into the non-canonical metabolism of *C. thermocellum*, factors limiting high product titer, and inform troubleshooting efforts aimed at engineering *C. thermocellum* for the production of various useful products (e.g. ethanol, 2,3-butanediol, etc.) at high titer.

As a cellulolytic specialist, *C. thermocellum* possesses a relatively narrow substrate range. This raises the question of whether substrate transport or metabolism (i.e., enzyme interconversion) is the primary barrier limiting the utilization of alternative carbon substrates in *C. thermocellum*. Sugar transport in *C. thermocellum* is generally not well understood, however, prior work has elucidated (Nataf et al. 2009) and reviewed (Tjo and Conway 2024) several ABC sugar transporters in *C. thermocellum*. In regards to substrate metabolism, *C. thermocellum* only ferments a small number of known substrates (i.e., cellulose, cellobiose, Avicel, etc.) (McBee 1954; Zhang and Lynd 2005; Lynd and Weimer 2002). *C. thermocellum* has also been adapted for growth on glucose, fructose, and sorbitol (Johnson et al. 1985; Nochur et al. 1990; Yayo et al. 2021; Daley et al. 2024) and engineered with a heterologous pathway to enable the consumption of xylose (Tafur Rangel et al. 2020), a pentose wood sugar that it cannot naturally grow on (Verbeke et al. 2017). However, the extent of substrate utilization in this organism has not been systematically studied.

Building on prior research, the goal of this study is to use alternative carbon substrates as a tool to advance understanding of the substrate range and non-canonical central metabolism of *C. thermocellum* to inform efforts aimed at engineering this microbe for high titer production of fuels and chemicals. Towards this goal, we aimed to experimentally address the following questions: What carbon substrates can be transported into *C. thermocellum*? Which alternative carbon substrates can be consumed (metabolized) by *C. thermocellum* and what are the resulting fermentation products? Finally, how do substrates with different degrees of reduction affect redox balance and the distribution of fermentation products in *C. thermocellum*?

## Materials and Methods

### Strains

This work was performed with three different strains of *C. thermocellum*: 1) Wild-type (WT) *C. thermocellum* (strain LL1004), 2) a strain of *C. thermocellum* previously engineered for high ethanol production (strain LL1570) (Hon et al. 2018), and 3) WT *C. thermocellum* (strain LL1004) adapted to grow on 10 g/L sorbitol plus 10 g/L pyruvate in the absence of cellobiose (strain LL1936). WT *Escherichia coli* (strain MG1655) was used as an aerobic control strain for performing tetrazolium dye assays using the BIOLOG PM1 and PM2A MicroPlates.

### Growth Medium and Batch Fermentation Conditions

All batch fermentations of *C. thermocellum* were performed in MTC-5 growth medium (Hogsett 1995) containing varying concentrations of substrate(s) (reported in g/L) in 50 mL screw-cap (Falcon) tubes (Corning Science, REF 352098) with a 20 mL or 30 mL medium volume or in 150 mL glass serum bottles with a medium volume of 20 mL. All cultures were grown at 55°C. The duration of time for each fermentation varied based on the experiment. Refer to the figure captions for additional experiment-specific details.

### Fermentation Sampling and Preparation for HPLC Analysis

A 700 µL fermentation sample was taken from each culture and transferred to a 2.0 mL microcentrifuge tube (USA Scientific; SealRite; Cat. No.: 1620-2700). The sample was centrifuged for 5 minutes at 21,300 rcf using a tabletop centrifuge to pellet the cells. 400 µL of supernatant was transferred to a new 2.0 mL microcentrifuge tube. The supernatant was acidified with 20 µL of a 10% w/v sulfuric acid solution prepared in MilliQ water. The sample was vortexed for 10-15 seconds, then the acidified supernatant was transferred into a spin filter tube (Costar Spin-X Centrifuge Tube Filter; 8160; 0.22 µm Cellulose Acetate filter) and centrifuged for 5 minutes at 21,300 rcf to filter the sample. Following centrifugation, the filter was discarded and the tube containing the filtrate was capped and placed in a 4°C freezer for up to 1-2 weeks. 200 µL of each filtered sample was transferred into a HPLC vial (Spectrum Chemical Mfg Corp; 842-22590-PK) and capped (ThermoScientific; SureSTART; 6ARC11RRT1C).

### Standard Preparation for HPLC Analysis

#### Mixed Metabolite Standards

Mixed metabolite HPLC standards containing cellobiose, pyruvate, malate, succinate, lactate, formate, acetate, MOPS, and ethanol (Table 1) were prepared at a 1x concentration in MTC-5 medium components (potassium citrate monohydrate (Fisher Chemical; Product No.: C6H7K3O8; CAS-No.: 6100-05-6), citric acid monohydrate (Sigma Aldrich; Product No.: C0706; CAS-No.: 5949-29-1), potassium phosphate monobasic (Fisher Chemical; Product No.: P380-212; CAS-No.: 7778-77-0), MOPS sodium salt (Sigma Aldrich; Product No.: M9381; CAS-No.: 71119-22-7), and MilliQ water) and diluted to prepare 2x and 8x diluted standards for use as a three-point standard curve for metabolite quantification. Following preparation, 500 µL aliquots of each standard were transferred into separate 2.0 mL microcentrifuge tubes and stored at - 20°C prior to use. A 400 µL aliquot of each 1x, 2x, and 8x standard was transferred into a new 2.0 mL microcentrifuge tube, acidified with 20 µL of a 10% w/v sulfuric acid solution, and vortexed for 10-15 seconds to mix the sample. A 200 µL aliquot of each acidified standard was transferred to two separate HPLC vials and run at the beginning and end of each batch of fermentation samples on the HPLC. A 40 µL injection volume was used.

**Table 1:**
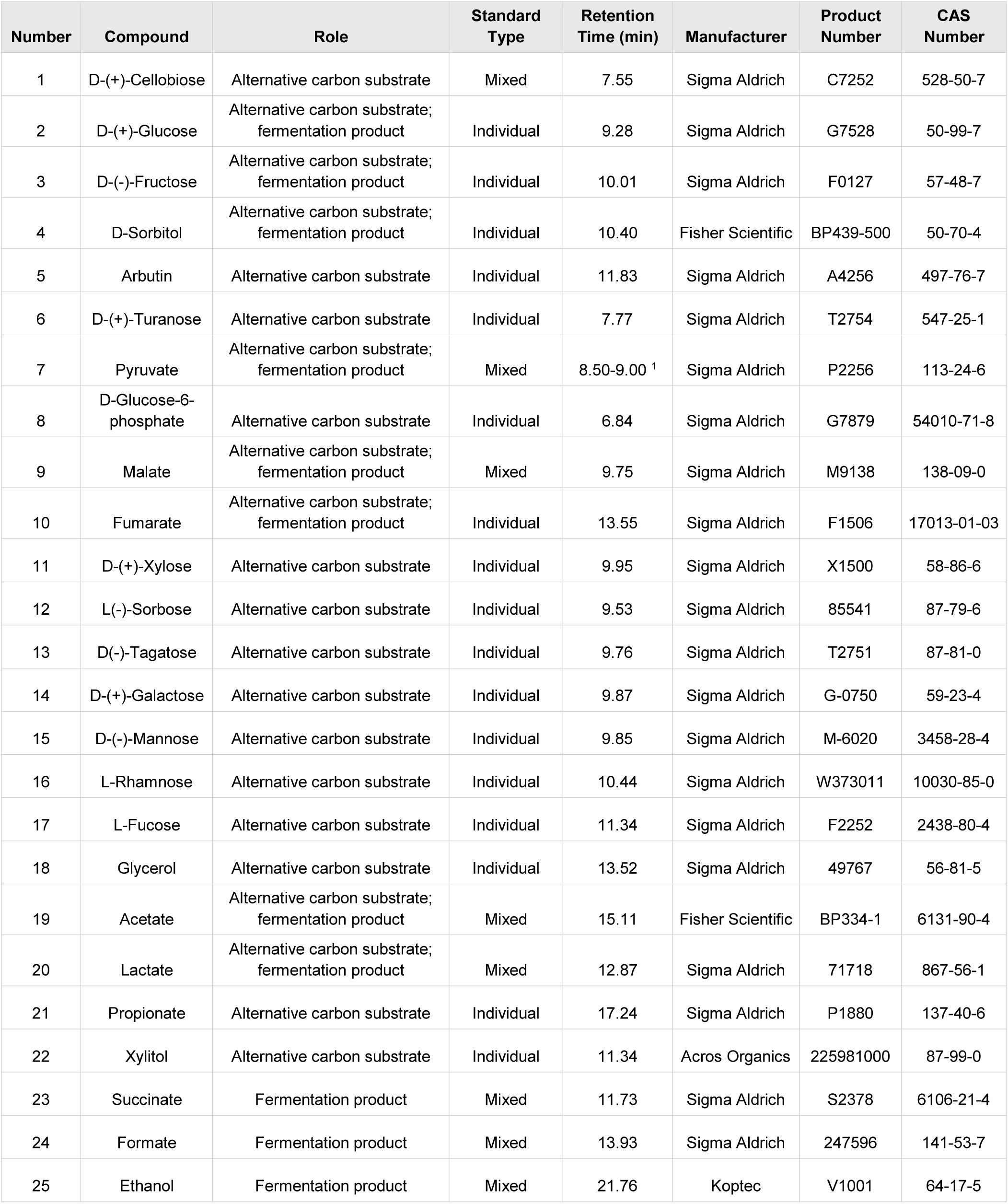
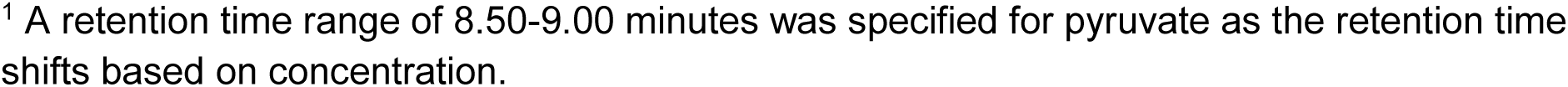
Alternative Substrates and Fermentation Products Examined in this Work.

| Number | Compound | Role | Standard Type | Retention Time (min) | Manufacturer | Product Number | CAS Number |
| --- | --- | --- | --- | --- | --- | --- | --- |
| 1 | D-(+)-Cellobiose | Alternative carbon substrate | Mixed | 7.55 | Sigma Aldrich | C7252 | 528-50-7 |
| 2 | D-(+)-Glucose | Alternative carbon substrate; fermentation product | Individual | 9.28 | Sigma Aldrich | G7528 | 50-99-7 |
| 3 | D-(-)-Fructose | Alternative carbon substrate; fermentation product | Individual | 10.01 | Sigma Aldrich | F0127 | 57-48-7 |
| 4 | D-Sorbitol | Alternative carbon substrate; fermentation product | Individual | 10.40 | Fisher Scientific | BP439-500 | 50-70-4 |
| 5 | Arbutin | Alternative carbon substrate | Individual | 11.83 | Sigma Aldrich | A4256 | 497-76-7 |
| 6 | D-(+)-Turanose | Alternative carbon substrate | Individual | 7.77 | Sigma Aldrich | T2754 | 547-25-1 |
| 7 | Pyruvate | Alternative carbon substrate; fermentation product | Mixed | 8.50-9.00 <sup>1</sup> | Sigma Aldrich | P2256 | 113-24-6 |
| 8 | D-Glucose-6-phosphate | Alternative carbon substrate | Individual | 6.84 | Sigma Aldrich | G7879 | 54010-71-8 |
| 9 | Malate | Alternative carbon substrate; fermentation product | Mixed | 9.75 | Sigma Aldrich | M9138 | 138-09-0 |
| 10 | Fumarate | Alternative carbon substrate; fermentation product | Individual | 13.55 | Sigma Aldrich | F1506 | 17013-01-03 |
| 11 | D-(+)-Xylose | Alternative carbon substrate | Individual | 9.95 | Sigma Aldrich | X1500 | 58-86-6 |
| 12 | L-(-)-Sorbose | Alternative carbon substrate | Individual | 9.53 | Sigma Aldrich | 85541 | 87-79-6 |
| 13 | D(-)-Tagatose | Alternative carbon substrate | Individual | 9.76 | Sigma Aldrich | T2751 | 87-81-0 |
| 14 | D-(+)-Galactose | Alternative carbon substrate | Individual | 9.87 | Sigma Aldrich | G-0750 | 59-23-4 |
| 15 | D-(-)-Mannose | Alternative carbon substrate | Individual | 9.85 | Sigma Aldrich | M-6020 | 3458-28-4 |
| 16 | L-Rhamnose | Alternative carbon substrate | Individual | 10.44 | Sigma Aldrich | W373011 | 10030-85-0 |
| 17 | L-Fucose | Alternative carbon substrate | Individual | 11.34 | Sigma Aldrich | F2252 | 2438-80-4 |
| 18 | Glycerol | Alternative carbon substrate | Individual | 13.52 | Sigma Aldrich | 49767 | 56-81-5 |
| 19 | Acetate | Alternative carbon substrate; fermentation product | Mixed | 15.11 | Fisher Scientific | BP334-1 | 6131-90-4 |
| 20 | Lactate | Alternative carbon substrate; fermentation product | Mixed | 12.87 | Sigma Aldrich | 71718 | 867-56-1 |
| 21 | Propionate | Alternative carbon substrate | Individual | 17.24 | Sigma Aldrich | P1880 | 137-40-6 |
| 22 | Xylitol | Alternative carbon substrate | Individual | 11.34 | Acros Organics | 225981000 | 87-99-0 |
| 23 | Succinate | Fermentation product | Mixed | 11.73 | Sigma Aldrich | S2378 | 6106-21-4 |
| 24 | Formate | Fermentation product | Mixed | 13.93 | Sigma Aldrich | 247596 | 141-53-7 |
| 25 | Ethanol | Fermentation product | Mixed | 21.76 | Koptec | V1001 | 64-17-5 |

<sup>1</sup> A retention time range of 8.50-9.00 minutes was specified for pyruvate as the retention time shifts based on concentration.

#### Individual Substrate Standards

Individual sets of standards were separately prepared for each of the alternative carbon substrates examined in this work (Table 1). Each standard was prepared at three different concentrations in MTC media components and 500 µL of each standard was aliquoted into 2.0 mL microcentrifuge tubes. These standards were stored, prepared, and run on the HPLC using the same method as the mixed metabolite standards described above.

### HPLC Analysis of Fermentation Samples and Standards

All fermentation samples and standards were run on a Shimadzu Prominence-i LC2030C 3D Liquid Chromatograph HPLC system equipped with a BioRad Organic Acids HPLC column (Bio-Rad Aminex HPX-87H Column; 300 x 7.8 mm; Cat. No: 1250140). A 2.5 mM sulfuric acid solution (140 µL of a 95.0%-98.0% sulfuric acid solution (Fisher Chemical; A300SI-212; CAS-No: 7664-93-9) added to 1.0 L of MilliQ water) was used as the mobile phase with a flow rate of 0.6 mL/minute. The runtime for each sample was 30 minutes. The samples were quantified using a refractive index detector (Shimadzu Corporation; Model: RID-20A; Cat. No.: 228-45104-42; Serial No.: L21465300941 CD) and a photodiode array (PDA) detector (integrated in the Shimadzu Prominence-i LC2030C 3D Liquid Chromatograph HPLC system). All fermentation samples were quantified in concentration units of millimolar (mM). Please refer to (D Richardson et al. 2025) for a detailed protocol regarding the maintenance, setup, method parameters, and data analysis used for running and analyzing all fermentation samples and standards on the HPLC reported in this work.

### Derivatization of Fermentation Samples for Amino Acid Quantification via HPLC

To quantify amino acids, a 40 µL aliquot of unacidified supernatant was derivatized using the methods, reagents, and standards provided in the Waters AccQ-Fluor Reagent Kit for Hydrolysate Amino Acid Analysis (REF: WAT052880; LOT: KIT0350481). The derivatized samples were subsequently run on the HPLC equipped with a Waters AccQ-Tag column (Waters AccQ-Tag Column For Hydrolysate Amino Acid Analysis; 3.9 x 150 mm; Part No.: WAT052885; Serial No.: 12123318612610) and three mobile phases: A) Waters AccQ-Tag acetyl-phosphate buffer (Waters AccQ-Tag Eluent A Concentrate for hydrolysate amino acid analysis; REF: WAT052890; LOT: 0152140261). This buffer was prepared by diluting 100 mL of eluent A concentrate into 1000 mL of sterile-filtered MilliQ water, B) 100% Acetonitrile (Sigma Aldrich; 34851-4L; CAS-No: 75-05-8), C) sterile-filtered MilliQ water. Each sample was run for 50 minutes using the gradient method provided in the kit instructions and quantified in concentration units of mM using a Shimadzu RF-20AXS fluorescence detector (AD1) (Shimadzu Corporation; Model: RF-20AXS; Cat. No.: 228-45148-42; Serial No.: L20505350205) (250 nm excitation, 395 nm emission). All derivatized samples were injected at 2 µL and 10 µL in case any amino acids oversaturated the Shimadzu RF-20AXS fluorescence detector. A powder standard containing all 20 natural amino acids provided in the Waters AccQ-Fluor Reagent Kit was derivatized and run on the HPLC using the same methods as the fermentation samples. The standard was injected at 1, 2, 4, 8, and 10 uL for the creation of a five-point standard curve.

### Tetrazolium Dye Assays

Tetrazolium dye assays (B R Bochner and Savageau 1977); (Barry R. Bochner et al. 2001); (Bhupathiraju et al. 1999) were performed using BIOLOG Phenotype MicroArray PM1 (BIOLOG; Product No.: 12111) and PM2A (BIOLOG; Product No.: 12112) 96-well MicroPlates. The MicroPlates arrived from BIOLOG pre-loaded with 95 different, dried carbon substrates in each plate, enabling the analysis of a total of 190 different substrates (well A1 in both plates was a no-substrate negative control). The tetrazolium dye assays were performed with WT *C. thermocellum* (strain LL1004) cells and cell lysate and WT *E. coli* (strain MG1655) cells as a control. The materials, reagents, and methods used to perform the tetrazolium dye assays were adapted from the manufacturer’s instructions for preparing PM MicroPlates using anaerobic bacteria and is published on protocols.io (D Richardson et al. 2024)5).

### Batch Fermentations on Alternative Carbon Substrates

Batch fermentations with WT *C. thermocellum* (strain LL1004) were performed in triplicate on 22 alternative carbon substrates in 50 mL screw-cap (Falcon) tubes containing 20 mL of growth medium with 10 g/L alternative carbon substrate and 1 g/L cellobiose (to help initiate growth). An inoculated (cells) and uninoculated (no cells) tube was prepared for each substrate. A stock solution was prepared in MilliQ water for each substrate with a final concentration ranging from 40 g/L - 100 g/L depending on the solubility limit of each substrate. Each substrate stock solution was sterile-filtered (Millipore Steriflip 50 mL Vacuum Filtration System with Millipore Express PLUS Membrane (0.22 µm pore size); Cat. No.: SCGP00525), and transferred into the anaerobic chamber to deoxygenate for 1-2 days prior to use. To each tube was added 10 mL of 2x growth medium (1 g/L cellobiose), sterile-filtered MilliQ water, and substrate stock solution to achieve a final concentration of 10 g/L alternative carbon substrate plus 1 g/L cellobiose with a final volume of 20 mL. 25 µL of strain LL1004 freezer stock was used to inoculate each tube.

### Batch Fermentations in Serum Bottles on 50 g/L Cellobiose, Fructose, or Sorbitol

#### Growth Medium and Bottle Preparation

Batch fermentations with *C. thermocellum* strain LL1570 were performed in triplicate in 20 mL of growth medium containing 50 g/L cellobiose, 50 g/L fructose, or 50 g/L sorbitol in 150 mL glass serum bottles. The glass serum bottles were capped, crimped and autoclaved prior to use. 100 mL stocks of 2x growth medium containing 50 g/L cellobiose, 50 g/L fructose, or 50 g/L sorbitol were prepared, sterile-filtered, and deoxygenated for 1-2 days prior to use. 20 mL of growth medium was added to each sterilized glass serum bottle using a 20 mL syringe (BD 20 ml syringe Leur-Lok Tip; REF: 302830) equipped with a 18 gauge needle (BD PrecisionGlide Needle; 18G x 1 ½” TW (1.2 mm x 40 mm); REF: 305196) and filter (Corning Sterile Syringe Filter; 0.20 µm pore size; Part No.: 431215). A 23 gauge needle (BD PrecisionGlide Needle; 23G x 1” TW (0.6 mm x 25 mm; REF: 305145) equipped with a filter was used to vent each bottle during growth medium addition. The bottles were nitrogen purged for 20 cycles at 45 seconds per cycle using pure nitrogen gas and stored at 4°C for 2 days prior to inoculation.

#### Inoculation, Sampling, and Pressure Measurements

A pre-culture of strain LL1570 was grown up in 30 mL of growth medium (5 g/L cellobiose) to an optical density (OD_600_) of 0.75. The glass serum bottles were transferred into an anaerobic glove bag, sterilized with 70% isopropyl alcohol, and punctured with a sterile 21G BD Precision Glide needle equipped with a filter for 15-20 seconds to remove any residual pressure in the headspace of the bottle. A 1.0 mL syringe equipped with a sterile 23G BD Precision Glide needle was used to inoculate each bottle with 700 µL of strain LL1570 pre-culture. All bottles were placed in a 55°C shaking incubator for 35 days. The pressure in the headspace of each bottle was measured periodically (in units of psi) using a pressure gauge equipped with a sterile, 23G BD Precision Glide needle. A fermentation sample was taken from each bottle on day 0 and 35 and periodically over the course of the fermentation using a 5 mL syringe equipped with an 18G needle, processed, and run on the HPLC using the methods described above.

#### Measurement of H_2_ Production

At the end of the fermentations, a 1.0 mL analytical syringe equipped with a needle (Vici Precision Sampling A-Series Pressure-Lok Analytical Syringe; Baton Rouge, LA; Part No.: 010033) was used to take a 1.0 mL sample from the headspace of each glass serum bottle. The sample was immediately injected onto a gas chromatography (GC) instrument (SRI Instruments 310C Gas Chromatograph) to measure hydrogen gas (H_2_) production.

### Batch Fermentations on Pyruvate with Increasing Concentrations of cellobiose

#### Overview and Reagent Preparation

Batch fermentations with WT *C. thermocellum* (strain LL1004) were performed in triplicate on 10 g/L sodium pyruvate plus 1 g/L, 2 g/L, 5 g/L, or 10 g/L cellobiose in 30 mL of growth medium in 50 mL screw-cap (Falcon) tubes. Batch fermentations were also performed on cellobiose only as a positive control. 500 mL of 2x growth medium containing 1 g/L, 2 g/L, 5 g/L, or 10 g/L cellobiose were separately prepared and deoxygenated for 1-2 days prior to use. A 40 mL 100 g/L sodium pyruvate stock solution was prepared in MilliQ water in a 50 mL screw-cap (Falcon) tube for each set of replicates, sterile-filtered, and deoxygenated for 1-2 days prior to use.

#### Fermentation Setup and Data Acquisition

The tubes for the cellobiose only conditions were prepared by adding 15 mL of 2x growth medium to 15 mL of sterile-filtered MilliQ water to achieve a final concentration of 1 g/L, 2 g/L, 5 g/L or 10 g/L cellobiose. The tubes for the sodium pyruvate and cellobiose conditions were prepared by adding 15 mL of 2x growth medium to 12 mL of sterile-filtered MilliQ water and 3 mL of 100 g/L sodium pyruvate stock solution to achieve a final concentration of 10 g/L sodium pyruvate plus 1 g/L, 2 g/L, 5 g/L or 10 g/L cellobiose. Three tubes were prepared for each condition: 1) a cellobiose only tube, 2) an inoculated (cells) pyruvate and cellobiose tube, and 3) an uninoculated (no cells) pyruvate and cellobiose tube to control for non-biological pyruvate degradation in growth medium. Each tube was inoculated with a 1% inoculum of strain LL1004 pre-culture grown to an optical density (OD_600_) value of 0.60. A 700 µL sample was taken periodically over the course of the fermentation, processed, and run on the HPLC using the methods described above. The optical density (OD_600_) of each culture was also measured using a spectrophotometer every 1-2 days during the fermentation. At the end of the fermentations, a 1.0 mL sample was taken from each tube, derivatized and run on the HPLC using the Waters AccQ-Tag HPLC column to quantify amino acids using the methods described above.

#### Measurement of Cell Carbon

The mass of the cell pellet in each 50 mL screw-cap (Falcon) tube was measured at the end of the fermentations for the purpose of calculating the cell carbon as a component of the reported carbon balances. For each culture, a new 50 mL screw-cap (Falcon) tube was placed in a 102°C oven to dry for 24 hours. The weight of the dry tube was measured three times and averaged. Each culture tube was centrifuged to pellet the cells and the supernatant was removed. The cell pellet was washed 3x in 25 mL of MilliQ water to remove any residual growth medium. Each pellet resuspended in MilliQ water was transferred to a dried and weighed 50 mL screw-cap (Falcon) tube. The cultures were centrifuged and the water was removed. All of the tubes were placed in a 102°C oven to dry for 24 hours. The weight of each tube containing the dried washed cell pellet was measured three times and averaged.

### Adaptation of WT *C. thermocellum* to Growth on a Mixture of Sorbitol and Pyruvate

#### Batch Fermentations on a Mixture of Sorbitol and Pyruvate

Batch fermentations of WT *C. thermocellum* (strain LL1004) were performed in triplicate in 30 mL of growth medium containing a mixture of 10 g/L sorbitol and 10 g/L sodium pyruvate in 50 mL screw-cap (Falcon) tubes. 500 mL of 2x growth medium (0 g/L cellobiose) and separate 100 g/L stock solutions of sorbitol and sodium pyruvate in MilliQ water were prepared, sterile-filtered, and deoxygenated for 1-2 days prior to use. 15 mL of 2x growth medium (0 g/L cellobiose), 9 mL of sterile-filtered MilliQ water, 3 mL of 100 g/L sorbitol stock solution, and 3 mL of 100 g/L sodium pyruvate stock solution was added to each tube. The tubes were inoculated with 100 µL of LL1004 pre-culture grown to an optical density (OD_600_) of 0.60. An inoculated and an uninoculated tube was prepared for each replicate.

#### Adaptation to Growth on a Mixture of Sorbitol and Pyruvate

To adapt strain LL1004 to growth on a mixture of 10 g/L sorbitol and 10 g/L sodium pyruvate, a 1% aliquot of spent growth medium from replicate 1 from the initial set of batch fermentations was inoculated into 30 mL of fresh growth medium containing 10 g/L sorbitol plus 10 g/L sodium pyruvate. This new culture was designated as subculture 1. The optical density (OD_600_) was measured and 700 µL samples for HPLC analysis were taken at the beginning, end, and every 1-2 days over the course of the fermentation of subculture 1. Upon reaching mid-exponential growth phase (OD_600_ = 0.6-0.8), a 1% aliquot of subculture 1 was inoculated into 30 mL of fresh growth medium containing 10 g/L sorbitol plus 10 g/L sodium pyruvate, which was designated as subculture 2. This subculturing process was repeated for a total of five subcultures. A freezer stock was prepared for each of the five subcultures upon reaching mid-exponential growth phase. The freezer stock for subculture 5 was added to the strain collection as strain LL1936, which is strain LL1004 adapted to growth on 10 g/L sorbitol plus 10 g/L sodium pyruvate.

#### Identification of Mutations Enabling Growth on a Mixture of Sorbitol and Pyruvate

To examine the mutations present in strain LL1936 (adapted for growth on sorbitol plus pyruvate) that enabled efficient growth on 10 g/L sorbitol plus 10 g/L sodium pyruvate compared to strain LL1004, 30 mL of growth medium containing 10 g/L sorbitol plus 10 g/L sodium pyruvate was inoculated with 25 µL of strain LL1936. Upon reaching mid-exponential growth phase, this culture was plated on a dilute plate containing a 1:1 ratio of autoclaved 1.6% agar (0.8% final concentration) solution and growth medium (10 g/L cellobiose). 20 individual colonies were picked and inoculated into 12 mL of growth medium containing 10 g/L sorbitol plus 10 g/L sodium pyruvate. The first 15 cultures to reach an OD_600_ of ∼0.6-0.8 were centrifuged, processed, and sent to a sequencing service provider (Plasmidsaurus) for whole genome sequencing along with a culture of WT *C. thermocellum* (strain LL1004) grown on 5 g/L cellobiose as a control. The sequencing results received from Plasmidsaurus were analyzed using Geneious Prime (Version 2026.0.2) to identify the mutations present in strain LL1936.

### Carbon Balance Calculations

#### Cumulative and Differential Carbon Balance Calculations

For the batch fermentations of strain LL1570 performed on 50 g/L cellobiose, 50 g/L fructose, or 50 g/L sorbitol, differential carbon balances were calculated for each sampling interval by determining the change in concentration of each metabolite relative to the preceding time point, converting each concentration change using nominal C3-equivalent (i.e. three-carbon backbone) carbon-accounting multipliers, and summing the resulting values. The differential carbon balance was then expressed as the interval-specific carbon difference normalized to the initial carbon-accounting basis, defined as the initial substrate amount expressed in nominal C3-equivalent units, and multiplied by 100. Cumulative carbon balances were calculated by sequentially summing the interval-specific carbon differences and normalizing the resulting cumulative carbon difference to the same initial carbon-accounting basis.

#### Carbon Balance Calculations for Batch Fermentations on Pyruvate and Cellobiose

For the batch fermentations of strain LL1004 performed on 10 g/L sodium pyruvate plus 1 g/L, 2 g/L, 5 g/L, or 10 g/L cellobiose, the amounts of pyruvate and cellobiose consumed were calculated from the changes in their measured concentrations, while the final measured concentrations of ethanol, acetate, lactate, formate, valine, and alanine were used as the fermentation product values. Estimated CO₂ was calculated from the products expected to be accompanied by CO₂ formation, using one mole of estimated CO₂ per mole of ethanol, acetate, or valine produced. Lactate, formate, and alanine were assigned no associated CO₂ in this calculation. All substrate, product, and estimated-CO₂ concentrations were converted from units of millimolar to millimolar carbon atoms (mM C atoms) by multiplying by the number of carbon atoms per molecule. Cell-pellet carbon was determined from the difference between the average mass of the dried tube containing the cell pellet and the average mass of the empty dried tube, assuming that carbon constituted 50% of the dried-cell mass; this carbon mass was divided by the atomic mass of carbon, converted to millimoles, and normalized to the volume of medium remaining in the tube to obtain a concentration in mM C atoms.

## Results

Tetrazolium dye assays were performed to experimentally assess the ability of WT *C. thermocellum* (strain LL1004) to transport and metabolize 190 different carbon substrates (Figure 1). There were 8 substrates identified as positive hits in the PM1 plate and 10 substrates identified as positive hits in the PM2A plate using strain LL1004 cells (Figure 1C). There were 6 substrates identified as positive hits in the PM1 plate and 9 substrates identified as positive hits in the PM2A plate using strain LL1004 lysate. A positive hit was indicated by the presence of purple color in the plates loaded with cells or lysate and no color in the corresponding well in the negative control (no cells) plate. Wells with a visibly darker purple color in the plates loaded with cells or lysate compared to the negative control (no cells) plate were also designated as a positive hit. There were 14 substrates identified as false positives in the negative control (no cells) PM1 plate and 13 substrates identified as false positives in the negative control (no cells) PM2A plate. A false positive was indicated by the presence of a colored well (either purple or a different color) in the negative control (no cells) plate and the presence of either color or no color in the plates loaded with cells or lysate. False positives likely arose in the negative control (no cells) plates due to the presence of hydrogen gas (H_2_) at low concentrations in the anaerobic chamber reacting with and reducing the BIOLOG Redox Dye Mix D present in the master mix of reagents loaded into the plates (Figure 1B). This hypothesis was supported by the manufacturer’s instructions, which stated that the plates can absorb hydrogen gas, which in turn can reduce the BIOLOG Redox Dye Mix D and confound the assay results when the PM MicroPlates are prepared in an anaerobic chamber. Several strategies were implemented to minimize exposure of the PM MicroPlates and assay reagents to hydrogen gas in an effort to reduce the number of false positives. First, the PM MicroPlates were deoxygenated in an anaerobic box equipped with an oxygen-absorbing anaerobic sachet outside of the anaerobic chamber. The PM MicroPlates were only briefly exposed to hydrogen gas in the anaerobic chamber for a few minutes while the plates were loaded, sealed, and returned to the anaerobic box for subsequent incubation at 55°C outside of the anaerobic chamber. Second, the BIOLOG Redox Dye Mix D was stored long-term aerobically at 4°C and was only briefly introduced into the anaerobic chamber to prepare the master mix. We also attempted to prepare the plates both anaerobically and aerobically with a larger inoculum volume, however the anaerobic approach did not reduce the number of false positives and the aerobic approach resulted in the cells not remaining viable, yielding inconclusive results.

**Figure 1:**
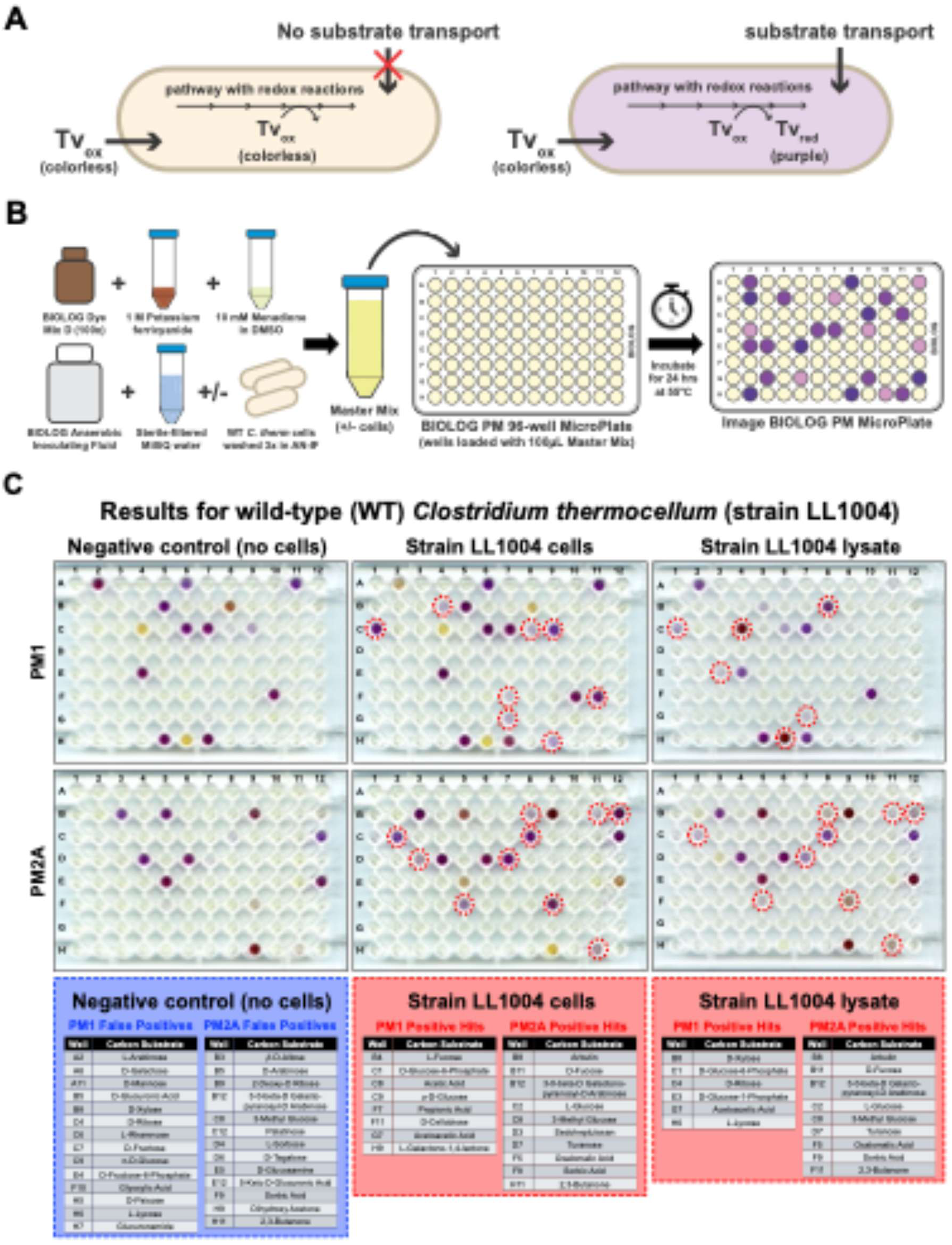
Tetrazolium dye assay for substrate uptake. **A)** Conceptual overview of the assay: when there is no substrate transport into the cell, the membrane-permeable tetrazolium dye remains oxidized (Tv_ox_) and colorless. When a substrate is transported into the cell, it is converted to various intermediate metabolites, some of which can reduce tetrazolium dye in the presence of certain metabolic enzymes. The reduced dye (Tv_red_) forms an insoluble purple compound that is visible by eye and absorbs strongly at 500 nm. **B)** Overview of the experimental method used to set up and run the tetrazolium dye assays using BIOLOG PM1 and PM2A MicroPlates. **C)** Results of the tetrazolium dye assays for WT *C. thermocellum* (strain LL1004). Wells with a red dashed circle represent positive hits, which indicate the transport and metabolism of the substrate. The tables below the plates provide a listing of the substrates that appeared as either false positives (blue box) in the negative control (no cells) plates or as positive hits (red boxes) in the plates loaded with strain LL1004 cells or cell lysate. The PM MicroPlates arrived from BIOLOG pre-loaded with a different dried substrate in each well. Refer to the supplementary data set for a listing of all the substrates present in the BIOLOG PM1 and PM2A MicroPlates (note that the concentrations of the substrates are kept secret by the manufacturer but are likely to be in the 1-10 g/L range). Refer to Figure S1 for an additional replicate of the tetrazolium dye assays results for strain LL1004. Refer to Figure S2 for the tetrazolium dye assays results for WT *Escherichia coli* (strain MG1655) performed as a control.

**Figure 2:**
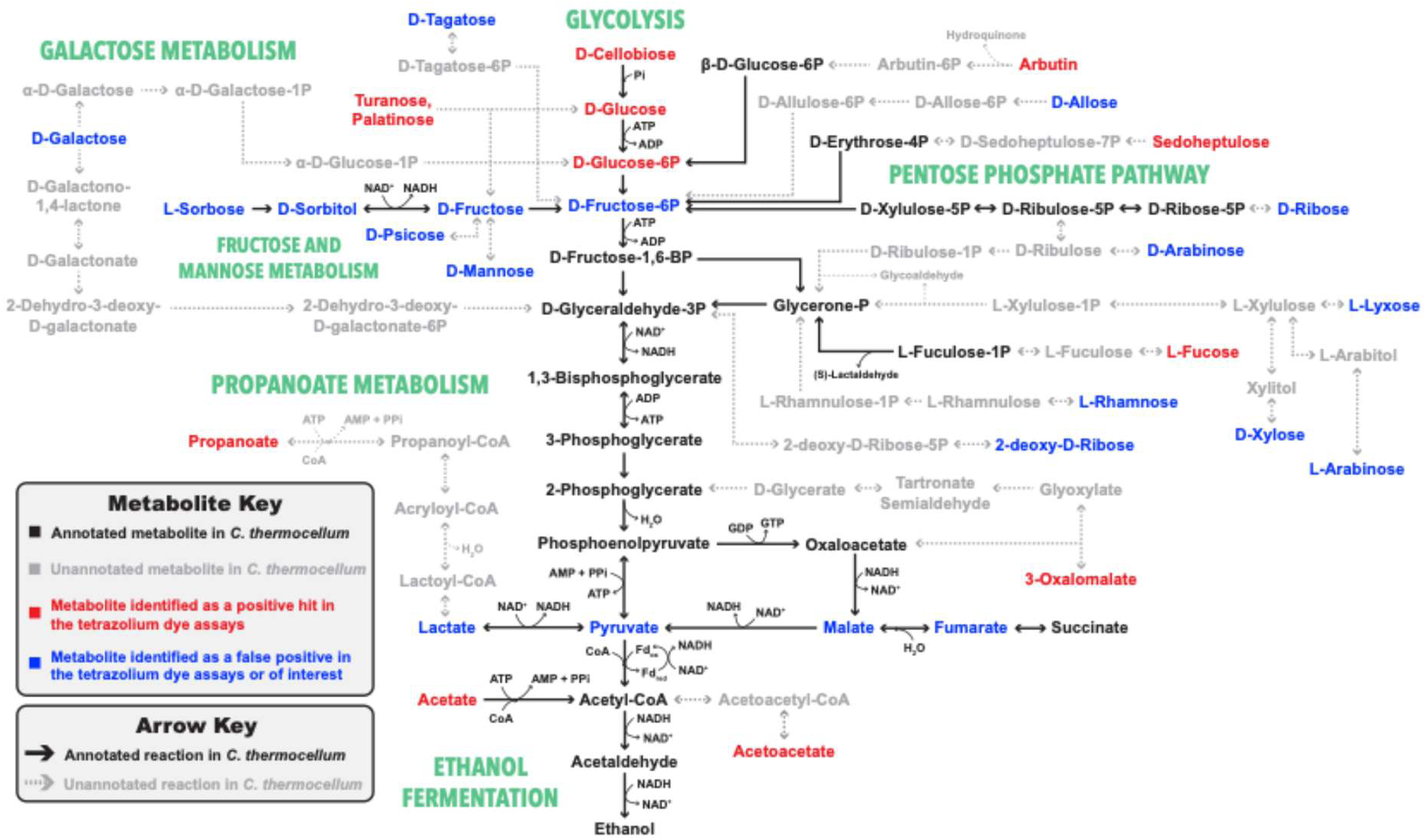
Map of central metabolism in WT *C. thermocellum* from cellobiose to ethanol. This map was developed using the Kyoto Encyclopedia of Genes and Genomes (KEGG) database for *Acetivibrio thermocellus* DSM1313 as a reference and includes metabolic pathways with annotated reactions (black solid lines) and pathways where the genes required for performing the reactions have not yet been annotated or do not exist in this microbe (grey dashed lines). The metabolites colored in red were substrates identified as positive hits in the tetrazolium dye assays. The metabolites colored in blue were substrates identified as false positives in the tetrazolium dye assays or were additional substrates of interest to study. All other metabolites were colored as black or gray based on if they were annotated or not in KEGG for *Acetivibrio thermocellus* DSM1313, respectively. For improved readability and simplicity, redox and energy cofactors, water, and coenzyme A (CoA) were not specified for all relevant reactions.

To further examine the ability of WT *C. thermocellum* (strain LL1004) to consume alternative carbon substrates, 22 substrates were selected for further experimental analysis. All of the substrates that were identified as positive hits and false positives in the tetrazolium dye assays were purchased for subsequent analysis if they were: 1) commercially available and 2) not prohibitively expensive (exceeding $500/g). In addition to cellobiose (1) as a control, the substrates identified as positive hits that were selected included: fucose (2), glucose-6-phosphate (G6P) (3), glucose (4), acetate (5), and propanoate (6) from PM1 and arbutin (7) and turanose (8) from PM2A. The substrates identified as false positives that were selected included: galactose (9), mannose (10), xylose (11), rhamnose (12) and fructose (13) from PM1 and sorbose (14) and tagatose (15) from PM2A. In terms of additional substrates of interest, sorbitol (16) was selected because: 1) it is more reduced than cellobiose in terms of its degree of reduction and may be able to help resolve redox imbalances, and 2) *C. thermocellum* has been previously adapted to grow on sorbitol (Johnson et al. 1985; Nochur et al. 1990; Daley et al. 2024). pyruvate (17) and glycerol (18) were selected as both substrates enter central metabolism at intermediate metabolites and have differing degrees of reduction relative to one another and cellobiose (pyruvate is less reduced and glycerol is more reduced than cellobiose). Malate (19) and fumarate (20) were selected as these metabolites participate in the incomplete tricarboxylic acid cycle in *C. thermocellum* and are converted to one another through a single chemical reaction (Xiong et al. 2018). We were curious to determine where the carbon from these substrates was directed through central metabolism. Lactate (21) was selected as it is a primary fermentation product of *C. thermocellum* when it is in a stressed state, such as high substrate loading or growth cessation (Thompson and Trinh 2017). Finally, xylitol (22) was selected as it is a 5-carbon sugar similar to xylose, and xylitol and xylose are interconverted through a single chemical reaction. To assess the extent of substrate consumption and product formation for each substrate, batch fermentations were performed with strain LL1004 on 10 g/L alternative substrate plus 1 g/L cellobiose (Figure 3).

**Figure 3:**
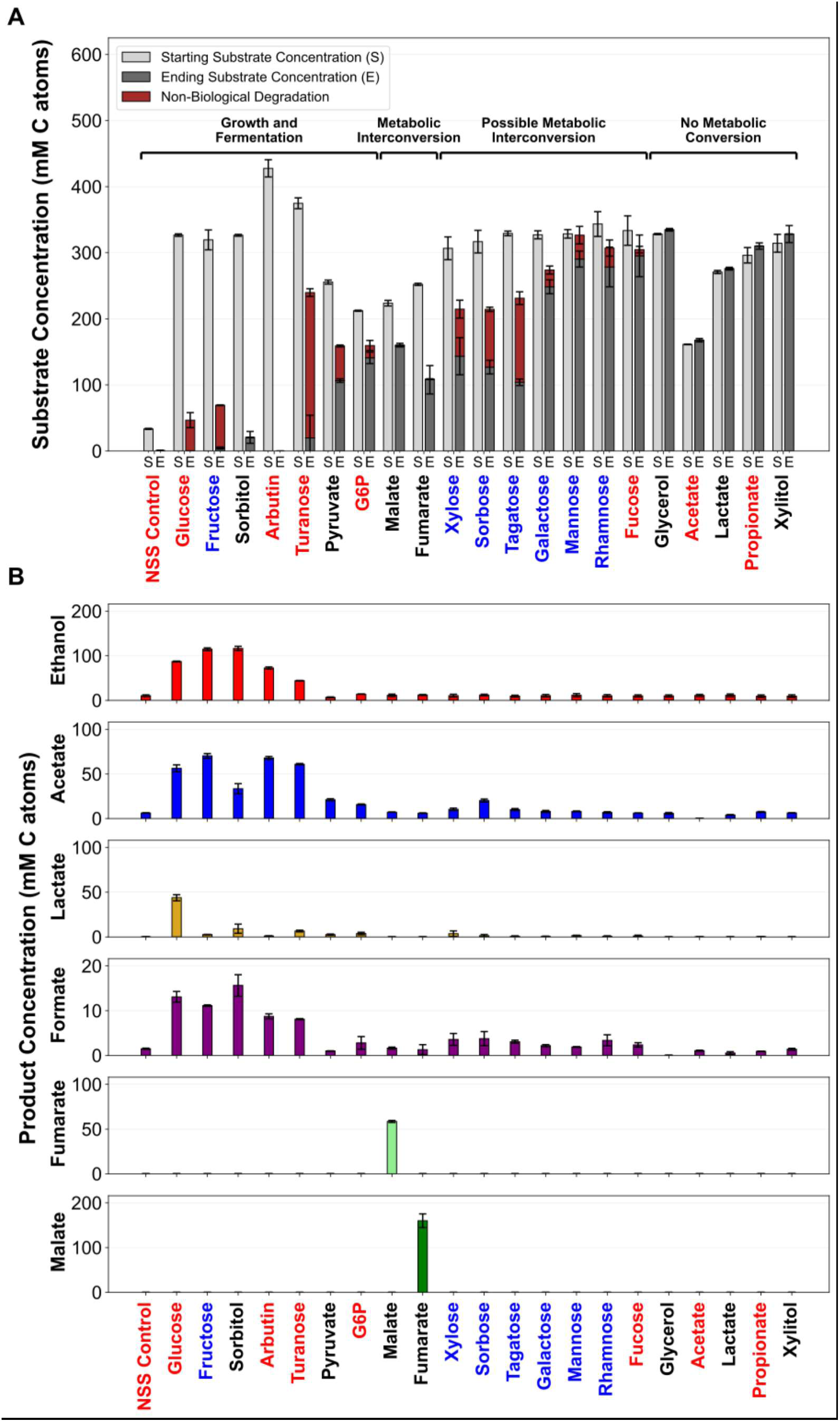
Substrate consumption and product formation for WT *C. thermocellum* (strain LL1004) grown on 22 alternative carbon substrates in 20 mL of growth medium containing 10 g/L alternative substrate plus 1 g/L cellobiose in 50 mL screw-cap (Falcon) tubes. Fermentations were run for 35 days. Samples were taken on days 0 and 35 and quantified by HPLC. **A)** Stacked bar plot showing the starting (S) and ending (E) concentrations of each substrate. Non-biological degradation was determined based on the decrease in substrate concentration in an uninoculated control. **B)** Fermentation products from each substrate. All results were quantified in units of mM C atoms. Error bars represent one standard deviation across three replicates. N = 3 independent fermentations. NSS Control stands for “no secondary substrate” control, which represents the 1 g/L cellobiose condition added to each substrate to help initiate growth. The x-axis substrate color indicates its results from the tetrazolium dye assays: red (positive hit), blue (false positive), black (negative hit). The underlying data for Figure 3 is provided in the supplementary data set.

To organize the analysis of the batch fermentation results, the 22 substrates were grouped into four categories based on the extent of substrate consumption and product formation (Figure 3A). The ‘Growth & Fermentation’ category consists of substrates that were either fully or partially consumed and converted to fermentation products beyond the products arising from the 1 g/L cellobiose. This category includes several known substrates of *C. thermocellum*, such as cellobiose and glucose, along with a few substrates that *C. thermocellum* has been previously adapted to grow on (i.e., glucose, fructose and sorbitol) (Johnson et al. 1985; Nochur et al. 1990; Yayo et al. 2021; Daley et al. 2024). A few additional substrates placed in this category were arbutin, turanose, pyruvate, and glucose-6-phosphate (G6P). The second category was ‘Metabolic Interconversion’, indicating products that showed evidence of biologically-mediated consumption, and corresponding production of some other product. Malate and fumarate were included in this category as these substrates were primarily converted to one another through a single reaction. ‘Possible Metabolic Interconversion’ was a third category to characterize the substrates that showed evidence of biologically-mediated consumption but were converted to unknown fermentation products. These substrates included xylose, sorbose, tagatose, galactose, mannose, rhamnose, and fucose. Glycerol, acetate, lactate, propionate, and xylitol were placed in the ‘No Metabolic Conversion’ category that includes compounds with no evidence of biologically-mediated conversion.

In terms of product formation, there was minimal fermentation product formation (i.e., production of ethanol, acetate, lactate, formate) outside of the substrates included in the ‘Growth and Fermentation’ category (Figure 3B). Of the substrates included in this category, the 1 g/L cellobiose condition served as a positive control to account for the products arising from the 1 g/L cellobiose in the other substrate conditions. Glucose was primarily converted to ethanol, acetate, lactate, and formate, and produced substantially more lactate than any of the other substrates. Fructose and sorbitol produced more ethanol than any of the other substrates. Growth on sorbitol produced considerably less acetate but more lactate and formate than growth on fructose. Arbutin (a glycosylated hydroquinone) was fully consumed and converted primarily to ethanol, acetate, and formate. A large, unidentified peak on the HPLC traces from the arbutin fermentations was consistent with disassociation of its hydroquinone ring. Growth on turanose produced less ethanol and acetate, more lactate, and roughly equimolar quantities of formate compared to growth on arbutin. Pyruvate was partially consumed by WT *C. thermocellum*, converted primarily to acetate, and produced minimal quantities of ethanol, lactate, and formate.

Several substrates including turanose, fructose, pyruvate, xylose, sorbose, and tagatose exhibited varying extents of non-biological degradation in the growth medium. The non-biological degradation of these substrates likely arose due to reaction with a component(s) in the growth medium.

Overall, many of the substrates that were identified as positive hits in the tetrazolium dye assays were either partially consumed or not consumed at all by WT *C. thermocellum* in this set of fermentations. It is also worth noting that different fermentation product profiles were observed for many of the substrates that were at least partially consumed, indicating that some of these substrates provide metabolic diversity beyond what is present in the canonical substrates fermented by this organism.

Following the batch fermentations on 22 different substrates, we decided to examine sorbitol as an alternative substrate in more detail. Based on our results with the WT strain (LL1004) showing that sorbitol shifted carbon flux toward ethanol and away from acetate (Figure 3B), we were interested to see if this effect could be used to increase ethanol titers in a strain of *C. thermocellum* engineered for high yield ethanol production (strain LL1570). To do this, we grew this strain on 50 g/L cellobiose, 50 g/L fructose, or 50 g/L sorbitol with no supplementation of cellobiose (Figure 4).

**Figure 4:**
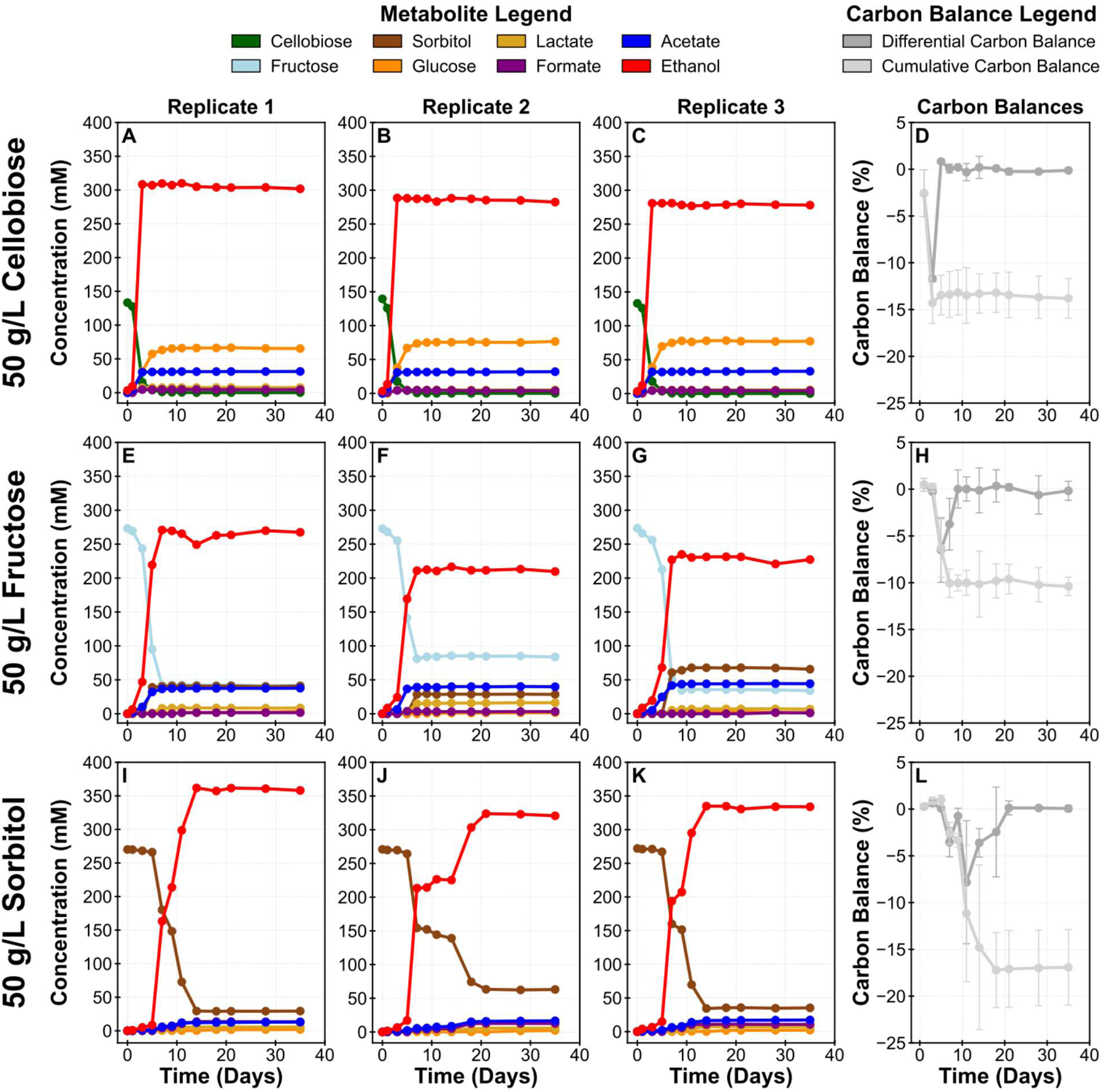
Time-course batch fermentations of *C. thermocellum* strain LL1570, previously engineered for high yield ethanol production (i.e. ethanologenic phenotype). Strain LL1570 was grown in 20 mL of growth medium containing 50 g/L cellobiose, fructose, or sorbitol in 150 mL glass serum bottles. **A-D)** Results for 50 g/L cellobiose. **E-H)** Results for 50 g/L fructose. **I-L)** Results for 50 g/L sorbitol. All fermentation products were quantified in units of millimolar (mM). The differential and cumulative carbon balances were quantified as a percentage difference from closed, where 0% represents a closed carbon balance. Differential carbon balances were measured as the difference of each pair of time points. Cumulative carbon balances were measured as the differences from time zero. Refer to the methods for a more detailed explanation of how the differential and cumulative carbon balances were calculated. Replicates are shown in separate panels, rather than as averages, due to lag-time variability. For panels D, H, and L, error bars represent one standard deviation across three replicates. N = 3 independent fermentations. Bottle pressure (measured in units of psi) was used as a readout for growth for these fermentations and the results are reported in Figure S3. Hydrogen gas (H_2_) production was measured at the end of the fermentations as is reported in Figure S4. The underlying data for Figure 4 is provided in the supplementary data set.

Of the three substrates, only the cellobiose was fully consumed. The fructose and sorbitol were only partially consumed. In terms of fermentation products, growth on 50 g/L sorbitol produced ∼300-350 mM ethanol, compared to ∼275-300 mM and ∼200-275 mM ethanol for 50 g/L cellobiose and 50 g/L fructose, respectively. In a similar manner to the WT strain, growth on 50 g/L sorbitol produced less acetate than growth on cellobiose or fructose. The 50 g/L sorbitol condition also produced more hydrogen gas (H_2_) than growth on 50 g/L cellobiose or 50 g/L fructose (Figure S4). We also observed an unexpected secondary lag phase across all three replicates of the 50 g/L sorbitol condition that resulted in a lag in the consumption of sorbitol and production of ethanol after about half of the substrate was consumed. To compare the differences between growth in glass serum bottles (with shaking during incubation) and growth in 50 mL screw-cap (Falcon) tubes (with no shaking during incubation), this set of fermentations was repeated in 50 mL tubes using strain LL1570 and similar results were obtained (Figure S5). These findings indicate that as a more reduced substrate than cellobiose and fructose, sorbitol may help to resolve redox imbalances and increase ethanol titers in strains of *C. thermocellum*.

Next, we wanted to understand how pyruvate affected metabolism. Pyruvate (C_3_H_4_O_3_ ^-^) is less reduced than cellobiose (C_12_H_22_O_11_) on a per-carbon basis, and we wanted to assess its effect on *C. thermocellum* metabolism compared to growth on sorbitol (a more reduced substrate). We chose to work with strain LL1004 (WT) to ensure maximal metabolic flexibility (i.e. no gene deletions which restrict metabolic flexibility, which are present in many engineered strains we have developed). Furthermore, To this end, we performed batch fermentations of WT *C. thermocellum* (strain LL1004) in 30 mL of growth medium containing 10 g/L alternative substrate plus 1 g/L, 2 g/L, 5 g/L, or 10 g/L cellobiose.

This fermentation provided several key insights into pyruvate metabolism in WT *C. thermocellum* (Figure 5). First, strain LL1004 can consume pyruvate in the presence of cellobiose, but it cannot consume pyruvate in the absence of cellobiose (Figure S8). Second, pyruvate inhibits growth. Third, pyruvate consumption is directly correlated with the amount of cellobiose present. As the concentration of cellobiose increases, the strain is less inhibited and it consumes more pyruvate. Fourth, the pyruvate plus cellobiose condition produced less ethanol and more acetate (center column) compared to growth on cellobiose only (left column). WT *C. thermocellum* produced valine and alanine and negligible quantities of the other natural amino acids across all of the pyruvate plus cellobiose conditions, which was absent in all of the cellobiose only conditions. Pyruvate stability in growth medium and milliQ water and the effect of pyruvate inhibition on the growth of strain LL1004 was further examined in Figures S9-S11.

**Figure 5:**
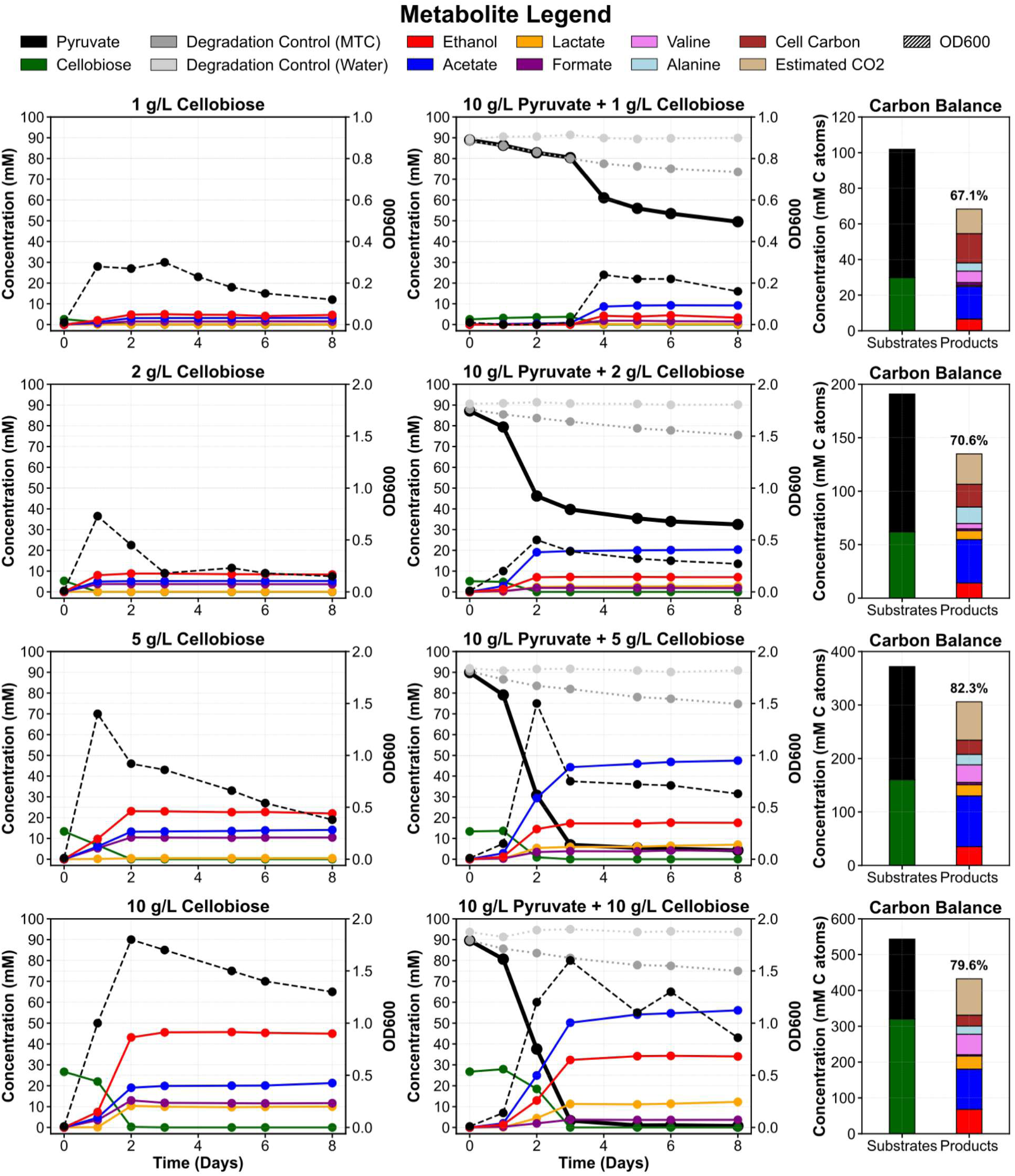
Time-course batch fermentations of WT *C. thermocellum* (strain LL1004) on pyruvate with varying concentrations of cellobiose. Fermentations were performed in 50 mL screw-cap tubes with 30 mL of growth medium containing 10 g/L sodium pyruvate plus 1, 2, 5, or 10 g/L cellobiose (center column). As a control, batch fermentations were also performed without pyruvate (left column). Carbon balances for the pyruvate plus cellobiose condition are provided in the right column. The black pyruvate bar within the substrates column of the carbon balances (right column) accounts for and excludes the non-biological degradation of pyruvate. The replicate 2 and 3 results for these fermentations are provided in Figures S6-S7. Refer to the methods for a description of how amino acids, estimated CO_2_, cell carbon, and carbon balances were calculated. The underlying data for Figure 5 is provided in the supplementary data set.

To further examine how feeding WT *C. thermocellum* (strain LL1004) with substrates with different degrees of reduction impacts redox balance and fermentation product profiles, we hypothesized that growth on a more reduced substrate (sorbitol) and a less reduced substrate (sodium pyruvate) than cellobiose could effectively cancel out in terms of its degree of reduction balance, and hypothesized that this would result in a fermentation product profile more similar to that on cellobiose. To test this hypothesis, we performed an initial set of batch fermentations of strain LL1004 on a mixture of 10 g/L sorbitol plus 10 g/L pyruvate in the absence of cellobiose (Figure S12). All three replicates of this fermentation took approximately 12-14 days to start growing, suggesting the possibility that mutation(s) may need to arise to enable growth on 10 g/L sorbitol and 10 g/L pyruvate in the absence of cellobiose (Figure S12A). These cultures simultaneously consumed the sorbitol and pyruvate and produced ethanol and acetate as the primary fermentation products (Figure S12B-D).

To adapt strain LL1004 to growth on a mixture of 10 g/L sorbitol and 10 g/L sodium pyruvate, a 1% aliquot of spent medium of replicate 1 (Figure S12) was inoculated into fresh medium containing 10 g/L sorbitol plus 10 g/L sodium pyruvate (subculture 1). Subculture 1 started growing on day 8 instead of day 12 (Figure 6). Upon reaching mid-exponential growth phase, a 1% aliquot from subculture 1 was transferred into fresh medium containing 10 g/L sorbitol plus 10 g/L sodium pyruvate. This process was repeated for a total of five subcultures and the optical density and fermentation product results are reported in Figure 6. Subculture 5 started growing within the first 24 hours of inoculation and is henceforth referred to as strain LL1936. These results demonstrate that we were able to successfully adapt WT *C. thermocellum* to growth on a mixture of 10 g/L sorbitol and 10 g/L sodium pyruvate in the absence of cellobiose.

**Figure 6:**
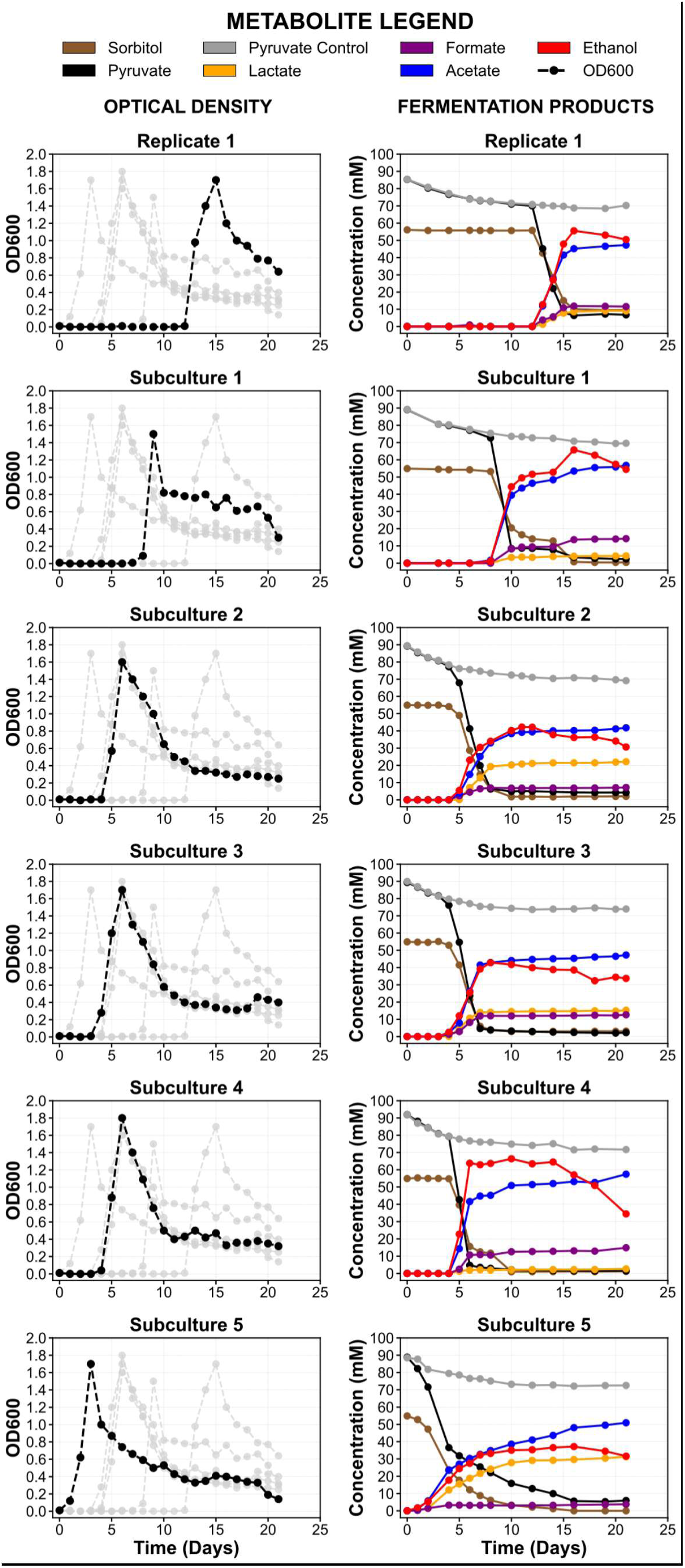
Adaptation of WT *C. thermocellum* (strain LL1004) to growth on a mixture of 10 g/L sorbitol and 10 g/L sodium pyruvate. Cells were transferred for a total of five subcultures. The adapted strain is referred to as LL1936. In the left column, the dashed black line represents the optical density of the specified subculture and the dashed grey lines represent the optical densities of the other subcultures. In the right column, the solid black line represents the pyruvate concentration and the solid grey line represents the pyruvate concentration in an un-inoculated control to account for its non-biological degradation in growth medium.

As the subcultures progressed, the adapted strain was found to primarily produce roughly equivalent concentrations of ethanol and acetate, and smaller quantities of formate. Lactate production was highly-variable with no obvious trend (Figure 6). Strain LL1936 was subsequently sent for full genome sequencing and the resulting table of mutations acquired from mapping its sequence to strain LL1004 (WT) grown on 5 g/L cellobiose as the reference genome is reported in Table S1.

## Discussion

Now we return to the original questions that motivated this work. The first was: what substrates is *C. thermocellum* able to transport and metabolize? Our initial approach involved using a tetrazolium dye assay to rapidly screen an existing library of 190 substrates (Biolog). A relatively small number (18/190) showed positive results (Figure 1C). Although we had initially suspected that this low number was due to substrate transport limitations, additional experiments with cell lysates showed similar results (15/190) (Figure 1C). Since the process of preparing the cell lysate removes the membrane as a barrier to transport, this contradicts the “transport limitation” hypothesis.

When we performed fermentations on these substrates (Figure 3), we chose a mix of both positive and negative hits from the tetrazolium dye assays. Furthermore, we had performed preliminary experiments showing that most of the BIOLOG carbon substrates do not support cell growth (supplementary data set), so we added 1 g/L cellobiose to identify substrates that could be co-fermented. Even in the presence of added cellobiose, only a handful of substrates showed any evidence of growth or metabolic conversion (Figure 3), demonstrating the limited substrate range of this organism. Interestingly, the tetrazolium dye assay turned out to be a relatively poor predictor of substrates that showed metabolic activity via the fermentation assays. For example: two of the most interesting substrates we studied in this manuscript (sorbitol and pyruvate) initially gave negative results from the tetrazolium dye assay, along with a few other substrates (malate and fumarate). We have several hypotheses to explain this discrepancy:

Our first hypothesis is that the difference is explained by the presence of a small amount (1 g/L) of cellobiose present in the co-fermentation assay that was absent from the dye assay. Cellobiose needed to be excluded from the dye assay, since that assay could not distinguish between metabolic activity associated with the initial cellobiose vs. activity associated with the target substrate. This explanation is most convincing for pyruvate, which is only fermented in co-fermentation experiments (i.e. with at least a small amount of initial cellobiose), and whose consumption increases linearly with the amount of initial cellobiose present (Figure 5). Thus, in the dye assay (with no cellobiose), there was no pyruvate-associated metabolic activity, and this activity only appeared in the presence of a co-substrate (e.g. cellobiose). The explanation is less convincing for sugars such as sorbitol and fructose, where growth was eventually observed without the need for a co-substrate (Figure 4).

A second hypothesis to explain the discrepancy between the co-fermentation and dye assays is the directionality of the dye assay. The idea behind using tetrazolium as an indicator of metabolic activity is that metabolism contains many redox reactions (some that oxidize redox cofactors (that would not likely interact with the tetrazolium dye) and some that reduce redox cofactors (that could give a positive dye result)), and that some of these will interact with the dye if metabolic activity is present. Dye interaction associated with the electron transport chain and mitochondria is well established (B R Bochner and Savageau 1977). Tetrazolium dyes are known to interact with ferredoxin-linked enzymes, and have been used for assays of ferredoxin:NAD^+^ oxidoreductases (Lo et al. 2017). However interactions with other areas of metabolism are less well-characterized, and this is further muddled by the presence of menadione (a redox mediator) in the Biolog dye mix that can allow additional redox interactions with other enzymes. One piece of evidence contradicting the directionality hypothesis is that both sorbitol and fructose gave negative results with the dye assay. Sorbitol and fructose can be interconverted via the enzyme sorbitol dehydrogenase. Sorbitol oxidation to fructose is associated with reduction of a cofactor (usually NAD^+^), and if the tetrazolium dye could accept electrons from this system, a color change could be observed. If directionality was the key factor in the dye assay, we would have expected sorbitol to yield a purple color, but not fructose. However we observed no color change with either.

A third hypothesis to explain the discrepancy between the co-fermentation and dye assays is genetic adaptation. The long duration (35 days) of the co-fermentation experiment provided a much longer time window than the dye assay (1 day), and the resulting metabolic activity observed in the co-fermentation experiments may have been a result of mutant strains that overtook the population. We have clear evidence for this in the case of sorbitol adaptation (Table S1) (Daley et al. 2024), and there are prior reports of mutations in *C. thermocellum* during fructose adaptation (Yayo et al. 2021).

The second major question that we set out to answer in this work is: how do substrates with different degrees of reduction affect metabolism? Although fermentation of sugars to ethanol is theoretically redox-balanced, existing strains of *C. thermocellum* that have been engineered for high-yield ethanol production (e.g. strain LL1570, (Hon et al. 2018)) still produce substantial amounts of non-desired products that are more-oxidized than ethanol (e.g. acetate), indicating that excess reducing equivalents are lost. We hypothesized that the extra reducing equivalents produced during growth on sorbitol would divert some of this carbon flux to ethanol, and our observations were in agreement with this expectation. Growth on sorbitol suppressed acetate production (as expected), some of which was diverted to increased ethanol production (also expected). Excess electrons from sorbitol to ethanol conversion were diverted to hydrogen production (Figure S4). Sorbitol fermentations exhibited lower carbon balances (vs. either cellobiose or fructose), suggesting that additional products are being produced that we did not measure. Importantly, our ability to perturb carbon balance based on the choice of substrate shows provides an approach that could be used in the future to better understand missing carbon in *C. thermocellum* fermentations, a longstanding problem for this organism (Ellis et al. 2012; Holwerda et al. 2012). One interesting carbon flux that has not been previously observed is the accumulation of sorbitol during fructose fermentations (Figure 4). Since this reaction acts as an electron sink, it likely explains both the increase in acetate production and decrease in ethanol titer relative to the cellobiose control. This is not likely a major source of carbon flux in WT *C. thermocellum* however, since the enzyme that converts fructose to fructose-6-phosphate (F6P) (fructokinase, PfkB (Daley et al. 2023)) is thermodynamically favored to run in the F6P-producing direction, and this aligns with our lack of observation of sorbitol production in cellobiose fermentations (Figure 4). It is also consistent with a lack of xylose (glucose) isomerase in this organism. This enzyme could enable the pathway glucose → fructose → sorbitol. However, this enzyme activity needed to be added by heterologous gene expression to enable xylose catabolism in *C. thermocellum* (Tafur Rangel et al. 2020), suggesting that it is not present in the WT strain.

The use of pyruvate as a co-substrate provided additional insight into *C. thermocellum* metabolism, but also raised some new questions. Since pyruvate has a lower degree of reduction (3.3 available electrons per carbon) vs. cellobiose (4 available electrons per carbon), we expected that growth on pyruvate would shift carbon flux away from ethanol and toward acetate, and this is in fact what we observed (Figure 5). *C. thermocellum* appears to have a fairly flexible metabolism that can accommodate substrates with different degrees of reduction by changing the ratio of fermentation products.

One question remains: what limits pyruvate uptake, transport or metabolism? In prior work, there are many examples of pyruvate accumulation during fermentation, so some system for pyruvate export from the cell must be present (Argyros et al. 2011; Holwerda et al. 2020, 2014). Nevertheless, we have never observed growth of *C. thermocellum* on pyruvate, and prior attempts at co-fermentation with pyruvate have been highly variable (unpublished results). In this work, we have finally identified conditions that allow co-fermentation of cellobiose and pyruvate (Figure 5), a phenotype that does not require any adaptation (suggesting that no adaptive mutations are needed). Our observations that pyruvate can be secreted into the medium during fermentation (Argyros et al. 2011; Holwerda et al. 2014, 2020), and that it can be taken up in the presence of a suitable co-substrate (such as cellobiose) suggests that transport does not limit pyruvate uptake by WT *C. thermocellum*.

If pyruvate is transported into *C. thermocellum*, what prevents this organism from using the substrate as a sole carbon source? Prior studies have established that 2-carbon compounds (e.g. ethanol) can be converted to upper glycolytic intermediates, such as glucose-6-phosphate, demonstrating that all of the reactions in glycolysis have at least some reversibility (Jacobson et al. 2020). Furthermore, conversion of pyruvate to acetate can generate energy (in the form of ATP), and provide reducing equivalents (reduced ferredoxin) via the pyruvate ferredoxin oxidoreductase reaction. One possible explanation is pyruvate toxicity. Pyruvate starts to inhibit growth around 10-20 g/L (Figure S11), and no growth was observed at pyruvate concentrations greater than 20 g/L (Figure S10). The mechanism of this inhibition is not known, however reverse flux through the pyruvate phosphate dikinase (PPDK) reaction would generate pyrophosphate (PPi), and PPi is known to regulate many enzymes in *C. thermocellum* metabolism (Daley et al. 2023; Kuil et al. 2023; Taillefer et al. 2015). Thus, perturbation of PPi levels via pyruvate uptake could dis-regulate these enzymes, blocking metabolism, however this is just one possible explanation among many.

Another interesting question raised by the pyruvate uptake experiment is the apparent stoichiometric ratio of pyruvate to cellobiose conversion. As cellobiose increases from 1 to 2 to 5 g/L (2.9 to 5.8 to 14.6 mM), pyruvate consumption increased from 22 to 40 to 75 mM (Figure 5). This is an approximate stoichiometric ratio of 7.5, 6.8 or 5.1:1 ratios. It is possible that all of the pyruvate was consumed at the 5 g/L cellobiose loading, artificially lowering that ratio, and suggesting that the 7.5:1 and 6.8:1 numbers may be more accurate. We do not have any clear candidates for the underlying metabolism that could create this stoichiometry, although since each cellobiose can be converted into four moles of pyruvate, the ratio of “pyruvate taken up from the medium” to “pyruvate generated from cellobiose” is in the range of around 2:1, which may be more biologically meaningful.

A final question related to the physiological effect of pyruvate on *C. thermocellum* physiology is whether the shift to more-reduced products (e.g. ethanol) observed during growth on sorbitol can be offset by co-fermentation with pyruvate. In theory, since one of these substrates is more-reduced than cellobiose (i.e. sorbitol) and one is less reduced (i.e. pyruvate), in the presence of both substrates, the cell should be able to control its redox balance via substrate choice (in addition to product profile, as usual). We observed that the cells still needed some time to adapt to the substrate mixture (Figure 6), and this lag phase was quite similar to the lag we observed when growing on sorbitol alone, suggesting that the mutations associated with growth in these conditions are related primarily to the adaptation for sorbitol consumption, and are not due to redox balance effects. Furthermore, we observed a roughly 1:1 ratio of ethanol to acetate during mid-log phase (Figure 6 - although this occasionally diverged in late stationary phase), which is less than the 4:1 ratio of ethanol to acetate observed in sorbitol fermentations (Figure S13), but more than the 1:2 ratio of ethanol to acetate observed in fermentations with pyruvate plus cellobiose (Figures 5, S6-S7).

Collectively, this work has advanced our understanding of the extent to which *C. thermocellum* can transport and metabolize alternative carbon substrates and has broadened our understanding of the factors that determine its relatively narrow substrate range. Despite an extensive search, we identified only a small number of additional substrates that this organism can transport and ferment, including: arbutin, turanose, pyruvate, and glucose-6-phosphate (G6P), confirming prior classification of this organism as a cellulose-fermenting specialist (Demain et al. 2005; Lynd and Weimer 2002). The results also provide support for using alternative carbon substrates that enter metabolism at different locations and possess different degrees of reduction as an experimental tool for perturbing metabolism, redox balance, and fermentation profiles to further our understanding of the non-canonical metabolism of anaerobic, thermophilic bacteria such as *C. thermocellum*. We showed that redox balance in *C. thermocellum* is relatively flexible, with acetate:ethanol ratios adjusting to accommodate substrates with different degrees of reduction. We also showed that ethanol titer in strains of C. thermocellum engineered for high yield ethanol production (i.e. strain LL1570) may be limited by electron availability. These are just a few of the new physiological studies enabled by the alternative substrates described in this report. We expect that future use of these substrates may unravel many longstanding mysteries in this organism, including those related to missing carbon and metabolic regulation.

## Acknowledgments

The authors would like to acknowledge Dr. Anthony (Ty) A. Lanahan, Ph.D. and Marybeth I. Maloney, M.S. for their assistance with ordering and receiving all of the materials and reagents used for conducting the experiments presented in this work. The authors would also like to acknowledge Dr. Lee R. Lynd, principal investigator of the Lynd Lab in the Thayer School of Engineering at Dartmouth College, for providing a shared laboratory space and equipment with the Olson Lab for conducting the research presented in this work.

## Author Contributions

**IDR:** Conceptualization, Data curation, Formal analysis, Investigation, Methodology, Validation, Visualization, Writing – Original Draft, Writing – Review & Editing

**BDS:** Conceptualization, Data curation, Formal analysis, Investigation, Methodology, Writing – Review & Editing

**EYM:** Data curation, Formal analysis, Investigation, Validation, Visualization, Writing – Original Draft, Writing – Review & Editing

**RSP:** Data curation, Formal analysis, Investigation, Visualization, Writing – Review & Editing

**DGO:** Conceptualization, Data curation, Formal analysis, Funding acquisition, Investigation, Methodology, Project administration, Resources, Supervision, Visualization, Writing – Review & Editing

---

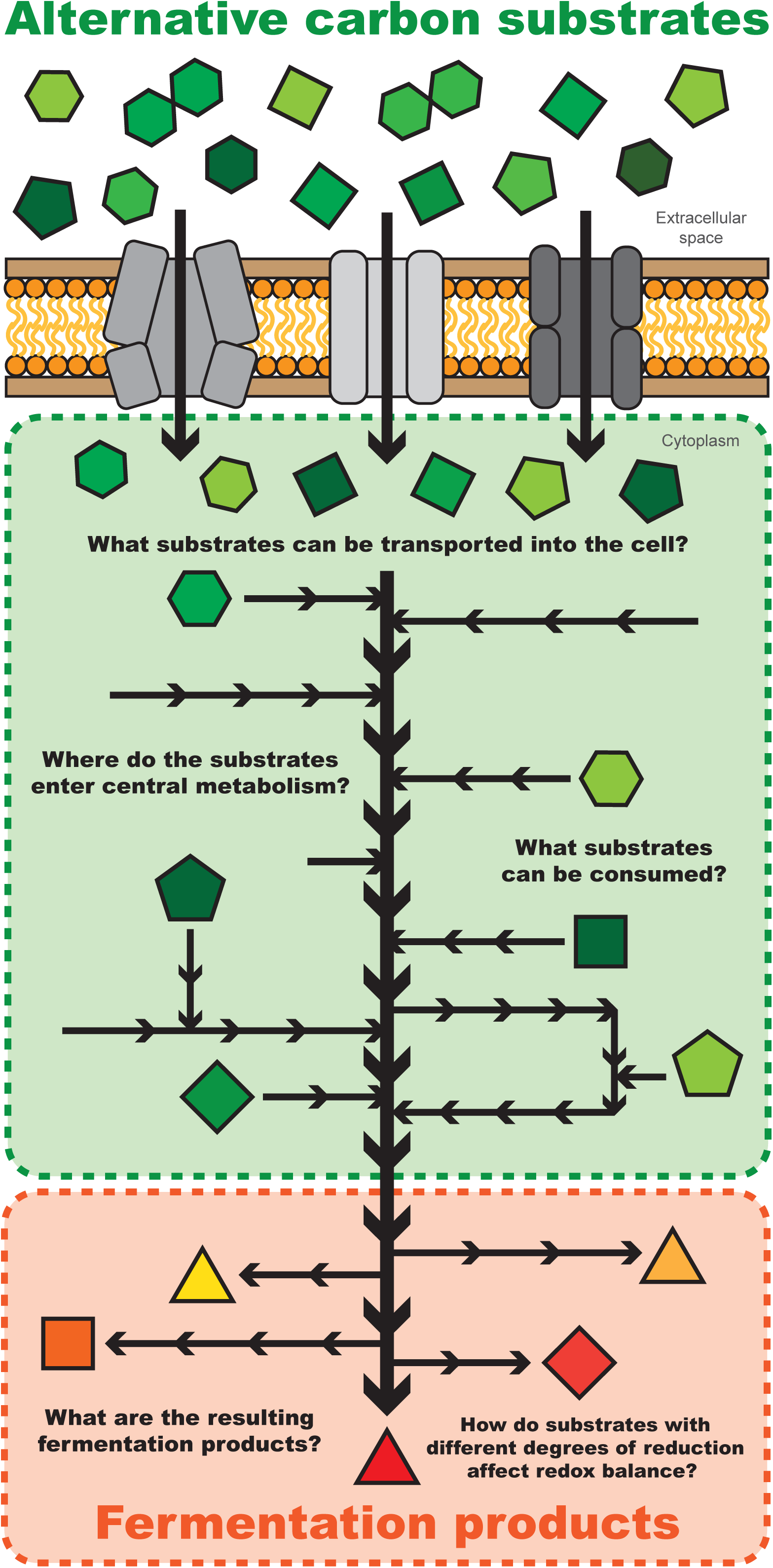

